# Social relationships among captive male pygmy slow lorises (*Nycticebus pygmaeus*): Is forming male same-sex pairs a feasible management strategy?

**DOI:** 10.1101/2020.10.01.318345

**Authors:** Yumi Yamanashi, Kei Nemoto, Josue Alejandro

**Author notes:** Correspondent author Yumi Yamanashi, Physical address: Okazaki kouen, Okazaki hosshoji-cho, Sakyo-ku, Kyoto City, Kyoto, Japan.

## Abstract

Little is known about the social behavior of pygmy slow lorises, in particular, the social relationships of same-sex individuals have rarely been investigated. The Slow Loris Conservation Center was built at the Japan Monkey Center to enhance the welfare of confiscated slow lorises, promote their conservation, improve public education and perform scientific research on the species. In the course of improving housing conditions, several same-sex pairs of pygmy slow lorises were formed. We monitored their behaviors and fecal glucocorticoid metabolite (FGM) levels to understand whether male same-sex pairings could be a feasible management strategy. The subjects were 10 male and 6 female lorises for comparison, all of whom were over five years old. We successfully formed five pairs of male lorises after eight formation attempts. Male pairs initially showed some aggressive behaviors; however, the rate decreased approximately 10 days after introduction. All of the male pairs eventually exhibited extensive affiliative social behaviors, including allogrooming and social play, during the dark (active) phase, and sleep site sharing during the light (inactive) phase. The rate of sleep site sharing during the light phase was higher than expected, suggesting that the pairs preferred to stay near each other. There was no evidence of increased stress after a long period of male–male social housing. Female same-sex pairs and male-female pairs demonstrated a high level of affiliative behaviors right after introduction. These results highlight the flexibility and high sociability of this species and indicate that such same-sex pairings are a feasible option for their social management.

## Introduction

Slow lorises (*Nycticebus spp*.) are in danger of extinction. The IUCN Red List currently classifies them into nine species, among which the pygmy slow loris (*Nycticebus pygmaeus*) is categorised as Endangered (IUCN, 2020). Although their international commercial trade has been prohibited as a general rule since 2007 by their inclusion in CITES Appendix I, one of the causes of their loss remains their frequent illegal trade (Musing, Suzuki, & Nekaris, 2015; Nekaris, 2007). Thus, an important conservation goal for this species is to reduce their use in the illegal pet trade through, among other strategies, public education (Norconk et al., 2020). Slow lorises are confiscated in both range and non-range countries, including Japan (Fuller, Eggen, Wirdateti, & Nekaris, 2018; Kitade & Naruse, 2020; Musing et al., 2015). Once confiscated, lorises are typically transferred to sanctuaries, rehabilitation centers or zoos (Fuller et al., 2018). Sanctuaries and rescue centers can often become strained past their capacity due to the unscheduled arrival of the confiscated animals (Moore, Wihermanto, & Nekaris, 2014), which can result in suboptimal captive conditions, mainly due to the lack of available space. Therefore, it is important to develop a practical strategy to house these individuals to ensure their well-being.

Little is known about the social behavior of wild pygmy slow lorises due to the inherent difficulty of nocturnal observation. The species was historically considered solitary, but studies have revealed that they are not asocial. Some slow loris species have been found to have spatial overlaps in home ranges between adult males and females (Wiens & Zitzmann, 2003), and friendly interactions between individuals have been reported (Wiens, 2002). Javan slow loris spend as much as 65% of their time in spatial proximity, including body contact for an average of 18% of their time (Nekaris, 2014). Studies have also demonstrated that male slow lorises are often found with scars, which has led some researchers to conclude that slow loris venom is used for intraspecific competition (Nekaris, Moore, Rode, & Fry, 2013; Nekaris et al., 2020). A recent study investigating the ranging patterns of pygmy slow loris in Cambodia found that the home ranges of some individuals overlapped with others to a small degree. Adult pygmy slow lorises display primarily solitary sleeping patterns, except for adult females with infants, who always sleep with their young (Starr & Nekaris, 2020). These solitary sleeping patterns are consistent with greater slow loris behavior (Wiens & Zitzmann, 2003), but different from Javan slow lorises, who often share sleep sites with conspecifics (Nekaris, 2014). Activity budgets differ across species and studies, with the proportions of social behaviors ranging from 0%–18% (Nekaris, 2014; Nekaris & Bearder, 2007; Rode-Margono, Nijman, Wirdateti., & Nekaris, 2014). Overall, the current available evidence suggests that slow lorises are more social than previously thought, and form some groups (Wiens & Zitzmann, 2003), including dispersed family groups (Javan slow loris and greater slow loris), and polygynous groups (pygmy slow loris) (Poindexter & Nekaris, 2020).

Slow lorises display social behaviors in captivity, not only between mother and infant (Fitch-Snyder & Ehrlich, 2003), but also between adults (Ehrlich & Alan, 1977). A previous study reported that two polygynous groups of greater slow lorises participated in various social behaviors, including allogrooming and social play (Ehrlich & Alan, 1977). Researchers believe that such stimulation from conspecifics can be a form of social enrichment (Fitch-Snyder, Schulze, & Larson, 2001). Despite these data, one survey focusing on the husbandry practices of North American zoos for lorisid primates revealed that approximately 55% of slow lorises are housed alone (Fuller, Kuhar, Dennis, & Lukas, 2013). Conspecific aggression during animal introductions was listed as one reason for the single housing. Although aggression can be a source of injury and stress in primates (Honess & Marin, 2006; Tennenhouse, Putman, Boisseau, & Brown, 2017; Yamanashi et al., 2016), and can compromise welfare if it continues over a long period or if the level of aggression is severe, moderate aggression is a natural behavior (Bloomsmith, 2001) that often occurs during the introduction phases of gregarious primates, and often decreases once their social relationships are established (Kutsukake et al., 2018). A well-established, cohesive social environment can offer ever-changing stimulation to animals in captivity. Successful social management can help to maximize the positive benefits of social groups and minimize negative effects, such as stress and aggression, but discussion on the practical social management of slow lorises is still limited.

Ideally, social structures in captivity should mimic the social structures seen in the wild for a particular species (Price & Stoinski, 2007); however, social structures in wild primates can vary in group size and composition depending on resource availability, predator presence, and other factors. In captivity, such environmental constraints differ (e.g. less resource competition and absent predators), which could allow for increased social group flexibility in captivity (Price & Stoinski, 2007). For example, same-sex groups are sometimes formed to control breeding or to manage surplus animals (Fàbregas & Guillén-Salazar, 2007; Sha, Alagappasamy, Chandran, Cho, & Guha, 2013; Stoinski, Kuhar, Lukas, & Maple, 2004; Stoinski, Lukas, Kuhar, & Maple, 2004; Yamanashi, Nogami, Teramoto, Morimura, & Hirata, 2018), even in species where such social compositions are non-existent in the wild (e.g. chimpanzees). Similarly, wild orangutans are semi-solitary species (van Schaik, 1999), but are sometimes kept in social groups in zoos (Amrein, Heistermann, & Weingrill, 2014; Weingrill, Willems, Zimmermann, Steinmetz, & Heistermann, 2011). Such flexible groups enable us to maximize space, and avoid suboptimal housing. However, the degree of social flexibility is species-dependent, and needs to be carefully investigated for each species. In slow lorises, same-sex pairs have also been formed in some captive settings (Fuller et al., 2013; Moore, Cabana, & Nekaris, 2015). In one study conducted at a rehabilitation center in Indonesia, confiscated slow lorises housed with conspecifics, particularly in same-sex pairs, displayed fewer stereotypic behaviors (Moore et al., 2015). However, the social relationships between same-sex slow lorises have rarely been investigated either in the wild or in captivity, and it is not well understood whether same-sex housing is a viable social management option for *Nycticebus spp*.

The purpose of this study was to investigate the social behaviors of male pairs of pygmy slow lorises, in order to consider whether same-sex pairing could be a feasible management strategy for this species. If slow lorises are motivated to interact with conspecifics, and benefit from being together, then we would consider that same-sex pairings are a feasible option. We particularly focused on male–male pairings in this study, because males often exhibit competitive tendencies (Van Schaik, 1994), which often makes their social management more complicated than female groups (Sha et al., 2013; Stoinski, Lukas, et al., 2004). In addition, we also measured the fecal glucocorticoid metabolite (FGM) levels as a measure of physiological stress. Glucocorticoids (GCs), the steroid hormones secreted by the adrenal glands of vertebrates, are frequently used as an indicator of physiological and psychological stress, because they are often increased when organisms face stressors (Selye, 1941; Squires, 2010). GCs generally increase available energy, and their acute increase is generally considered adaptive in stressful situations. However, prolonged exposure to GCs can result in a number of maladaptive consequences, such as neuronal cell death, insulin resistance, muscle and bone atrophy, poor wound healing, hypertension, and even immune system collapse to the point of death (Reeder & Kramer, 2005). Therefore, prolonged GC activation can be considered maladaptive from the view of animal welfare. For the purposes of this study, we explored five variables. First, we investigated the types of social relationships observed between adult males. Second, we determined how often sleep sites were shared, in order to understand whether the lorises preferred to stay together during their light phases (i.e. when the individuals are typically resting or sleeping). To exclude the potential influences of environmental factors, such as locational preference or temperature, we also recorded temperatures, and calculated the chance rate of sleep site sharing. Third, we observed the process that occurred during the formation of the social relationships, and its effects on the individuals’ physiological stress levels. Fourth, we compared the social relationship formation process across pairs of different sex combinations. Fifth, we considered the stability of each social relationship by comparing social behaviors observed during two different study periods.

If the pygmy slow lorises were motivated to interact with conspecifics, then we hypothesized that male pygmy slow lorises would display affiliative social behaviors, and prefer to share sleep sites during their light phases. In this case, physiological stress levels may decrease after social relationships have been formed, or they may remain the same. If slow lorises were not motivated to interact with conspecifics, then we hypothesized that they would choose to keep distance between each other. In the worst case, they would continue fighting without showing any sign of affiliation. Physiological stress levels in this case would then likely increase over time as a cost of the social environment.

## Methods

### Subjects

The subjects of this research were 16 pygmy slow lorises living at the Japan Monkey Center (JMC), in Inuyama, Aichi, Japan (10 males and 6 males). Fifteen of the individuals were confiscated between 2006 and 2007, and the final male was born at JMC. The confiscated slow lorises came from the customs office of airports in Japan. Specific information on each subject is given in Supplementary Table 1. Prior to the establishment of the Slow Loris Conservation Center (SLCC) at JMC, the subjects were housed individually in small cages due to space limitations (0.1–0.3 m^3^), except for one father-son pair. Once the center was established, the lorises were introduced to larger enclosures, as described below. This study was carried out in accordance with the recommendations in the “Guide for Animal Research Ethics” of the Japan Monkey Center, and the legal requirements of the country, and ASP Principles for Ethical Treatment of Non-Human Primates. The study protocol was approved by the institutional committee of the Japan Monkey Center (#2015001, #2016001, #2017003).

### Slow loris conservation center

The SLCC was established to improve the living conditions of confiscated pygmy slow lorises, and to serve as a place for environmental education, and slow loris research. The first slow lorises started to live there in 2015, and its facility and function gradually improved after that. The facility had previously been used for nocturnal primates, and was renovated by keepers and researchers to become the SLCC. Animal welfare is the first priority at SLCC, thus the renovations were carried out to match the needs of slow lorises. The facility uses a reverse-lighting cycle. Lighting levels were measured on 19 December 2019, and were found to be between 0.00 and 0.18 lux during the dark phase, depending on the distance from the light in each room. We used red-filmed light in each room during dark phase observations, in order to minimize the effects of lighting on the slow lorises (Ariana, Marco, & Nekaris, 2020; Fuller et al., 2016; Starr, Nekaris, & Leung, 2012). Enclosure size varied from 2.0 to 16.0 m^3^. For the male–male pair experiments, we used rooms that were about 8.0 m^3^. Branches were installed to facilitate locomotion, and nest boxes and hiding places were added to each enclosure (Fitch-Snyder, Schulze, & Streicher, 2008). The number of nest boxes was equal to, or more than, the number of individuals in the enclosure. The animals also used the branches and the floor for resting and sleeping. Diets mainly comprised of gum, insects and vegetables were provided in multiple locations in each enclosure (Cabana & Nekaris, 2015; Cabana & Plowman, 2014). The mean temperature was 25.8 ± 2.08 °C (measured between January and December 2017). Organized tours with a limited number of zoo visitors were held several times a month. Participants of the organized tour could explore the center using red light with educational staff, who described the ecology and conservation of slow lorises and the activities of JMC. Before arriving at the SLCC, most lorises lived alone except for the father-son pair (Yashi-Hiiragi). They were kept alone at least since 2013. The details of the social management in the past were not well-recorded, but some individuals had social housing experience. Based on husbandry records, some individuals confiscated on the same date were kept in the same cages for a while after their arrival, but they were separated because of aggression or death of some individuals. Although there is a possibility that some individuals were housed together for a few months up to a few years when they were young, Yanagi and Shuro were never housed together before the current study.

Social housing that restricted breeding was necessary to accommodate all the slow lorises housed at the facility when it was first established.

### Social introduction

Each pair was formed in an enclosure that were unfamiliar to both individuals. When necessary, we set up acclimation phases by placing small cages in close proximity for several days or by allowing the individuals to spend time in adjacent cages. In the beginning of this study, we conducted social introduction on a trial-and-error basis before establishing the described acclimation protocol. We judged whether the pairing could continue based on the behaviors exhibited. If the individuals immediately displayed aggressive behavior, and little to no affiliative behaviors were observed, we decided to stop the pairing and change the combination. If some affiliative behaviors were observed with few or no aggressive behaviors, or if no social behaviors were observed at all, then we continued the pairing. With this protocol, we formed five male–male pairs from eight formation attempts with nine individuals (one male formed two pairs with different individuals during different periods of the study, Supplementary Table 2). A tenth male always behaved aggressively; therefore, he was kept solitary at all times, and was not included further in the study. During the study of male-male pairing, they only had a visual access to females. The males lived in adjacent cages where they could visually, olfactorily, auditorily, and, possibly, tactilely socialize with other pairs. The females lived in cages a short distance away, with glass partitions in-between (Supplementary Figure 1).

### Data collection

The behaviors were recorded using infrared cameras (Zcam-CV, Bohankobo). The cameras were attached to the ceiling and recorded each enclosure. In most cases, it was difficult to identify the individuals from the recorded videos due to the video’s resolution, therefore, we recorded aggressive and non-aggressive interactions for the lorises without identifying each individual. An ethogram is provided in Table 1. We recorded aggressive behaviors using all-occurrence sampling methods (Martin & Bateson, 2007). We counted the bouts of aggressive behaviors. We included contact aggression in the category of aggressive interactions, as it is often difficult to differentiate non-contact aggression (i.e. chasing) from non-aggressive behaviors (i.e. following behaviors to inspect conspecifics) on the recorded videos. We recorded affiliative social behaviors that involved direct social contact (allogrooming, social play, co-feeding, nest box sharing) every 5 minutes during the dark phase. We categorized the affiliative behaviors based on (Fitch-Snyder & Ehrlich, 2003). We often observed that the two lorises entered a nest box, and categorized such behaviors as a form of affiliative social behavior. We sometimes observed the animals grooming each other or playing socially inside of the nest box, but could not observe all the behaviors that occurred inside the boxes. In addition to monitoring dark phase behaviors, we determined whether the lorises used the same sleep sites during their light phase. We primarily did so using recorded videos, but also used long-term data from Tsuge–Poplar, which were obtained by keepers (including the second author (KN)) during the light phase.

**Table 1.** Ethogram used for the study

| Category | Behaviors | Definition |
| --- | --- | --- |
| Affiliative behaviors | Social grooming | Lick fur and apply toothcomb while grasping other animal with one or both hands. |
|  | Play | Mock fight (mutual biting, hitting, clasping but no other signs of threat; low intensity, prolonged, neither animal tries to get away) |
|  | Co-feeding | Eat foods from the same dish in proximity |
|  | Sharing a nest box | Enter into a same nest box during dark phase |
|  | Sharing a sleeping site | Enter the same nest box or sleep in a close proximity, within reaching distance, during light phase |
| Aggressive behaviors | Attack | Bite, hit hard with hands or push with apparent force; lunge and bite at the same time, butt with head with apparent force |
|  | Fight | Mutual hitting and biting with apparent force; wrestle vigorously |

### Behavioral data recording

Behavioral data were collected between July 2016 and January 2020. Five types of behavioral collections were made. We used the data obtained in different periods for each purpose.

#### 1. Description of social behavior in five pairs of male slow lorises

Behaviors were recorded from the 5 pairs of male slow lorises (9 individuals) for a total of 150 hours between November 2016 and August 2017. We recorded their behaviors 5 hours a day from the start of the dark phase for 5 days, 3–4 months after the start of the social housing for each pair. We recorded the behavior for another 5 hours for 5 days and 1 year after the start of social housing for the Olive–Kunugi pair because the pair took a longer time to exhibit affiliative behaviors.

#### 2. Did they choose to stay in the same sleeping site during light phase?

Subjects were 5 pairs of male slow lorises (N = 9). A total of 450 days of sleeping site data were obtained between August 2016 and October 2019. Sleeping occurred primarily in their nest boxes, but the ground and branches were sometimes also used. We considered that slow lorises shared a sleeping site when they slept in close proximity, within reaching distance. Room temperature was recorded daily by keepers. To check whether sleep sites were shared by chance, we used sleeping site data collected over a month for each pair during the period when they formed affiliative relationships (four pairs, except for the father–son pair, for which data were collected for 16 days only because recordings were only available for this time period). To determine whether a relationship between sleep site sharing and temperature existed, we used data from three of the pairs. We selected the three pairs based on our ability to collect sleep site sharing data and room temperature data simultaneously for these three pairs during a period when they performed affiliative behaviors with almost no aggressive behaviors in their light phase. We collected data from the Tsuge–Poplar pair for 281 days, the Nagi–Poplar pair for 56 days and the Olive–Kunugi pair for 66 days.

#### 3. Process of social group formation and changes in fecal glucocorticoid metabolites in two pairs of male pygmy slow lorises

For this part of the experiment, we selected two pairs of male slow lorises as subjects (N = 4). We recorded their behaviors 5 hours a day from the day after introduction every day for 20 days, with additional sporadic observations after this period, for a total of 260 hours between August 2016 and August 2017. We started each observation from the start of the dark phase. Since the social relationships were established over different time periods for the different pairs, the timing of the observations varied.

#### 4. Sex differences in forming social relationships

The subjects were two pairs of male–male, female–female, and male–female pairs (N = 10). Two males experienced two types of pairings during different periods of the study. Total observation time was 144 hours between August 2016 and January 2020. We observed each female–female and male–female pair for 12 days for 2 hours from the start of the dark phase each day. Observations were made for the first 6 days after social housing began, and then for another 6 days one month after the start of social housing. Any injury incidents that occurred were recorded by keepers between August 2016 and January 2020.

#### 5. Stability of social relationship

Subjects were 3 pairs of male lorises (N = 6). Behaviors were compared between two periods; after 3–4 months, and after 8–12 months from the start of the social housing. We recorded their behaviors 3 hours each day from the start of the dark phase, for six days per observation period. The total observation time was 108 hours between November 2016 and August 2017.

### Fecal glucocorticoid metabolite (FGM) analysis

The feces from two pairs of male lorises were collected by keepers (N = 99) at 16:00 to 17:30 in the evenings (beginning of light phase). Fecal samples were collected 1–15 days before the start of social housing (11–14 days of sample collections for each pair) and during 0–31 days after the start of social housing (20–32 days of sample collections for each pair), between 15 July and 15 September, 2016. We could not identify the individual feces once the pairs were housed together. We cannot exclude the possibility of urine contamination, but the amount of slow loris urine is very small. Therefore, compared with large-bodied primates, urine contamination is less likely to occur. Before starting the social housings, individual feces were processed separately, and the average value was used for the analyses. From one day after social housing, all the feces collected were mixed and processed together. Collected feces were stored in freezers (−20°C) until use.

We extracted FGM based on the methodologies developed by Kinoshita et al. (2011). Feces were placed in a vacuum oven for about 18 hours at 60°C. The dried samples were pulverized using a hammer and contaminants were removed as much as possible using mesh. Approximately 0.02 g of the powdered samples were placed in a conical tube and 3 ml of 80% methanol was added to each tube. The tubes were shaken for 30 mins at ambient temperature, and then centrifuged for 10 mins at 3000 rpm. The supernatant was then placed into new 2.0 ml tubes, and dried in a vacuum oven at 60°C. The dried samples were reconstituted with the assay buffer before an assay. Cortisol levels were measured using enzyme immunoassay kits (Salimetric salivary cortisol kits: #1-3002). Parallelism was confirmed, and the recovery rates were 85.6% on average. Inter-assay variabilities were 3.92% (high) and 12.4% (low), respectively, and intra-assay variabilities were 4.41% on average. Data with CV values that were higher than 10% were removed from the analysis.

### Statistical analysis

We used R 3.6.1 to perform all the statistical tests described below (R Development Core Team, 2019). The data that support the findings of this study are openly available in Open Science Framework (Yamanashi et al., 2020).

### Did they choose to stay in the same sleeping site during light phase?

We performed two analyses, in order to determine whether the pairs chose to share sleep sites during the light phase. First, we compared the rate of sleep site sharing with the rate that could have occurred by chance. The chance level was calculated based on Cohen’s kappa calculation (Martin & Bateson, 2007). For example, suppose the two individuals, A and B, used two locations, *x* and *y*, for sleeping, and that data was obtained for 30 days. Individual A used sleeping location *x* a total of 11 times, and *y* a total of 19 times. Individual B used sleeping location *x* a total of 20 times, and *y* a total of 10 times. The chance proportion of sharing the location *x* can be calculated by: 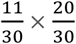 and that of sharing the location *y* can be calculated by: 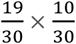. The chance proportion of sharing the same sleeping site can then be calculated as 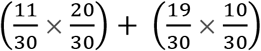. We calculated the chance proportion for each pair. Then, we checked whether the observed proportion of sharing the sleep sites was higher than the chance proportion calculated as above by binomial test. We used the function ‘pbinom’ in R.

We also checked the relationship between room temperature and sleep site sharing using Man–Whitney’s U tests for comparing temperatures when sleep sites were shared, and when they did not. We used the function ‘wilcox.exact’ of the package ‘exactRankTests’ to perform the analyses (Hothorn, Hornik, van de Wiel, & Zeileis, 2006).

### Changes in FGM levels

Fecal samples could not be assigned to specific individuals once the pairs were housed together. We compared the FGM levels among three conditions: 15 days before social housing (‘before’), 15 days right after social housing began (‘soon after’, 0–14 days) and days 15–31 (‘later’) after social housing began for each pair. For statistical analyses, we calculated the average values from each individual each day before social housing. We used the values obtained from FGM measurements of mixed samples after social housings. Among the four subjects, Nagi lived in a small cage and Poplar, Olive and Kunugi each lived in one of the enclosures (8.0 m^3^) during the ‘before’ period. We used Kruskal–Wallis tests to compare the FGM levels across the three conditions. If we statistically significant differences were determined, then we used Man–Whitney U tests for multiple comparisons with the Bonferroni correction.

### Sex differences in forming social relationships

The effect of sex on the formation process of social relationships was compared using a generalized linear model. We used the function ‘glm’ with Poisson distribution in R. Independent variables included sex (male–male, male–female, female–female), introduction phase (0 months or 1 month), pairs and the interaction of sex and introduction phase. Dependent variables included the number of times we observed aggressive and the number of sampling points we observed affiliative behaviors. We also looked for differences in the rate of affiliative behaviors one month after the start of social housing, by constructing a model that included the independent variables of sex and pair. We compared Akaike Information Criteria (AIC) among models composed of various combinations of the aforementioned factors to identify the models that produced the lowest AIC values. We used the dredge function in the package ‘MuMIn’ to choose the best models (Barton & Barton, 2019). The AIC tables, including the models containing ⊿AIC < 2 from the final model and the parameter estimates for the best models, are given in the supplementary materials.

### Stability of social relationships

We used a Mann–Whitney U test to compare the rates of social behaviors between two periods for each pair. We used a proportion test to compare the probability of sharing the sleeping sites.

## Results

### 1. Social behaviors observed in the five male–male pairs

All five pairs performed some affiliative behaviors. On average, the pairs continued for 595 ± 334 days (mean of four pairs), except for the father-son pair, who had lived together from the birth to the death of the son (Supplementary Table 2). Allogrooming, nest box sharing during dark phase and sleep site sharing during light phase were observed in all pairs (Table 2, Figure 1). Social play was observed in three of the pairs (Nagi-Poplar, Shuro-Yanagi, Yashi-Hiiragi: Supplementary video). Unlike other pairs, one pair (Olive–Kunugi) did not show any affiliative behaviors during the observations made three months after the start of the social housing; however, they did show affiliative behaviors similar to the other pairs after one year had passed.

**Table 2.** The rate of social behaviors among five pairs of male slow lorises (5 hours for 5 days/column) We recorded their behaviors 3–4 months after the start of the social housing for each pair and 1 year after the start of social housing for the Olive–Kunugi.

|  | Yashi–<br>Hiiragi | Shuro -<br>Yanagi | Nagi–Poplar | Olive–Kunugi |  | Tsuge–Poplar |
| --- | --- | --- | --- | --- | --- | --- |
| Sampling<br>period<br>(months) | Father-son | 4 | 4 | 3 | 12 | 3 |
| Aggression | 0 | 0 | 0 | 0 | 0 | 0 |
| Affiliation | 0.38 ± 0.043 | 0.14 ± 0.059 | 0.31 ± 0.14 | 0 | 0.27 ± 0.15 | 0.11 ± 0.084 |
| Sleep site<br>share(days) | 5 | 5 | 3 | 0 | 4 | 5 |

**Figure 1.**
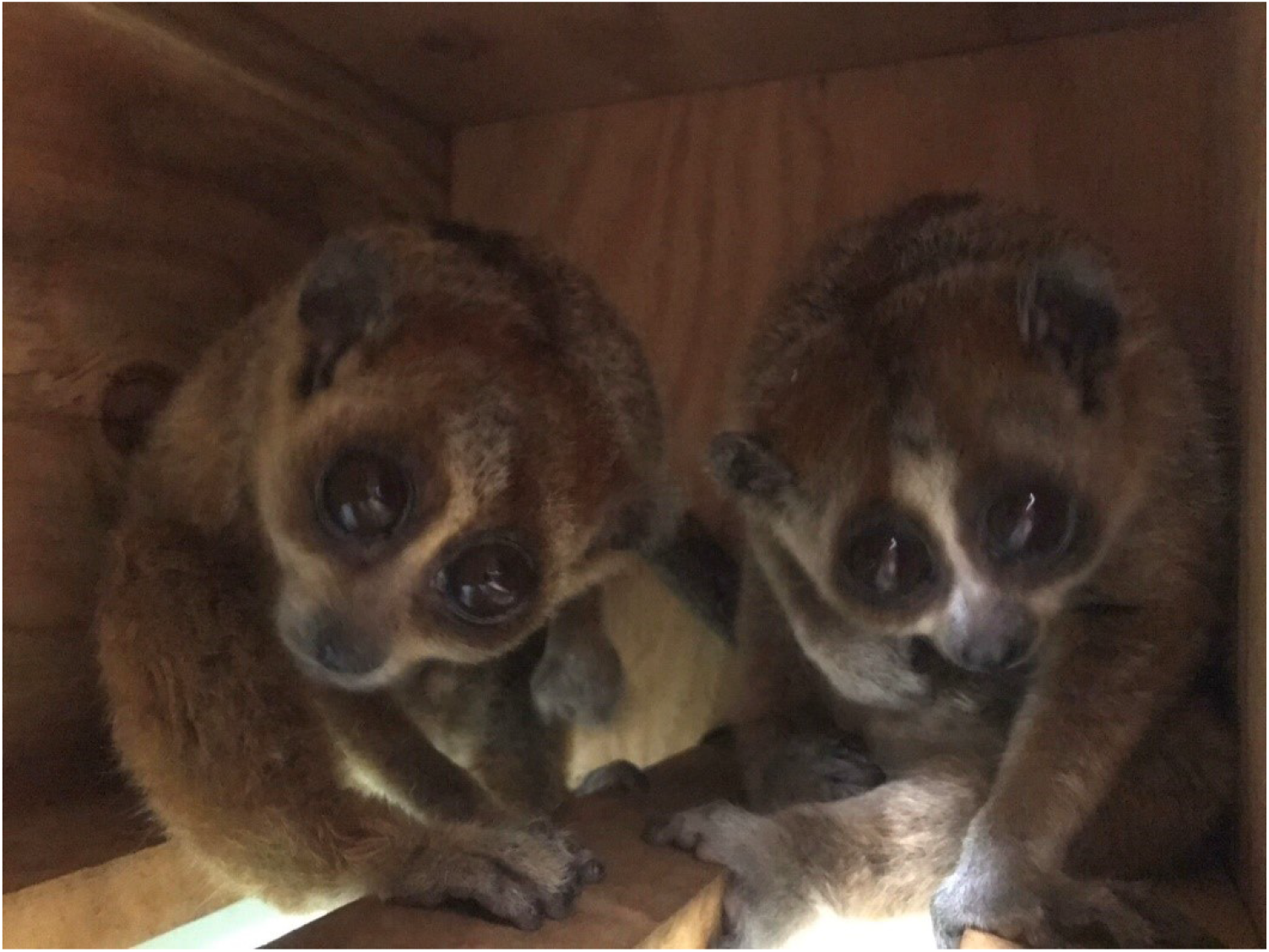
A photo of two males (Shuro and Yanagi) sharing a nest box

### 2. Did they choose to stay in the same sleeping site during light phase?

All pairs were observed to remain at the same sleeping site during the light phase. They used multiple locations for sleeping together. The proportion of sleep site sharing was above the calculated chance levels for all the pairs (Table 3).

**Table 3.** The proportion of sleep site sharing in the five male pairs

|  | Number of<br>sleeping site<br>used | Rate of sleeping<br>site share | Rate of sleeping site<br>share expected by chance | P |
| --- | --- | --- | --- | --- |
| Nagi-Poplar | 3 | 0.68 | 0.38 | <0.001 |
| Shuro-Yanagi | 2 | 1.00 | 0.85 | <0.001 |
| Olive-Kunugi | 3 | 0.81 | 0.68 | 0.018 |
| Tsuge-Poplar | 4 | 0.90 | 0.46 | <0.001 |
| Yashi-Hiiragi | 3 | 0.94 | 0.64 | <0.001 |

There was no statistically significant relationship between temperature and sleep site sharing for two of the pairs. The proportion of sleep site sharing was high for all the three pairs (Tsuge–Poplar: 96.4%; Nagi–Poplar: 67.9%; Olive–Kunugi: 84.8%). Nagi– Poplar had an average temperature of (mean ± SD): 26.2°C ± 0.98°C when they shared a sleeping site, and an average temperature of 26.3°C ± 0.73°C when they slept alone (Man–Whitney’s U test: W = 328, P = 0.81). Olive–Kunugi had an average temperature of 25.4°C ± 0.75°C when they shared a sleeping site, and an average temperature of 25.7°C ± 0.75°C when they did not (Man–Whitney’s U test: W = 218, P = 0.25). In the case of Tsuge–Poplar, the average temperature was relatively high when they shared sleep sites with an average temperature 24.7 °C ± 0.73 °C, and an average temperature of 24.2°C ± 0.67°C when they slept alone (Man–Whitney’s U test: W = 817, P = 0.029). It should be noted that they did not share sleep sites for 10 days out of 281, and that the temperatures when they did not share sleep sites were within the range of those when they shared.

### 3. Process of social group formation and changes in FGM in two pairs of male slow lorises

The male–male pairs showed some forms of aggression in the beginning of the study that ceased after about 10 days. One pair (Nagi–Poplar) gradually increased their affiliative behaviors after 14 days. It took more time for another pair (Olive–Kunugi) to exhibit comparable levels of affiliative social behaviors (Figure 2). Sleep site sharing during the light phase was observed after affiliative behaviors were observed during the dark phase. Aggressive behaviors were observed most often after intense following of another individual, during which the follower grabbed the other male and a short jostling occurred.

**Figure 2.**
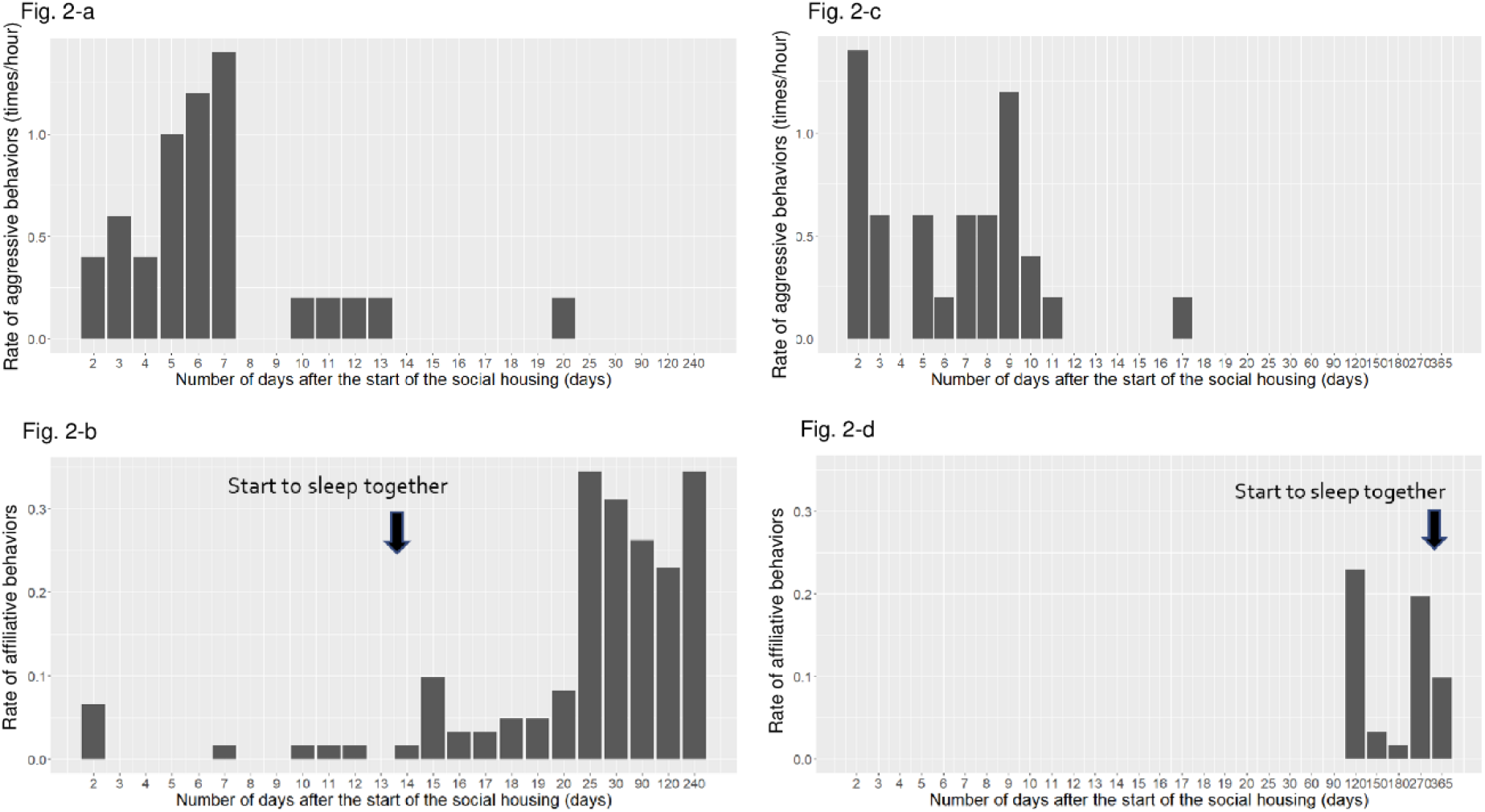
Changes in the social behavior from the start of the social housing in two pairs. Figure 2-a. Changes in aggressive behaviors for Nagi–Poplar. Figure 2-b. Changes in affiliative behaviors for Nagi–Poplar. Figure 2-c. Changes in aggressive behaviors for Olive–Kunugi Figure 2-d. Changes in affiliative behaviors for Olive–Kunugi

Changes in FGM levels are shown in Figure 3. In the case of Nagi–Poplar, there was a significant increase in FGM levels soon after the start of the experiment (Man– Whitney’s U test: ‘before’ vs ‘soon after’: W = 22, P < 0.001), but levels decreased to lower than those observed before the social housing began (Man–Whitney’s U test: ‘soon after’ vs ‘later’: W = 227, P < 0.001; ‘before’ vs ‘later’: W = 242, P < 0.001). There was no statistically significant change in FGM levels for Olive–Kunugi (Kruskal–Wallis test: X^2^ = 2.5, df = 2, P = 0.28). Before social housing, the FGM levels of Nagi were higher than those of Poplar (W = 164, P = 0.0091).

**Figure 3.**
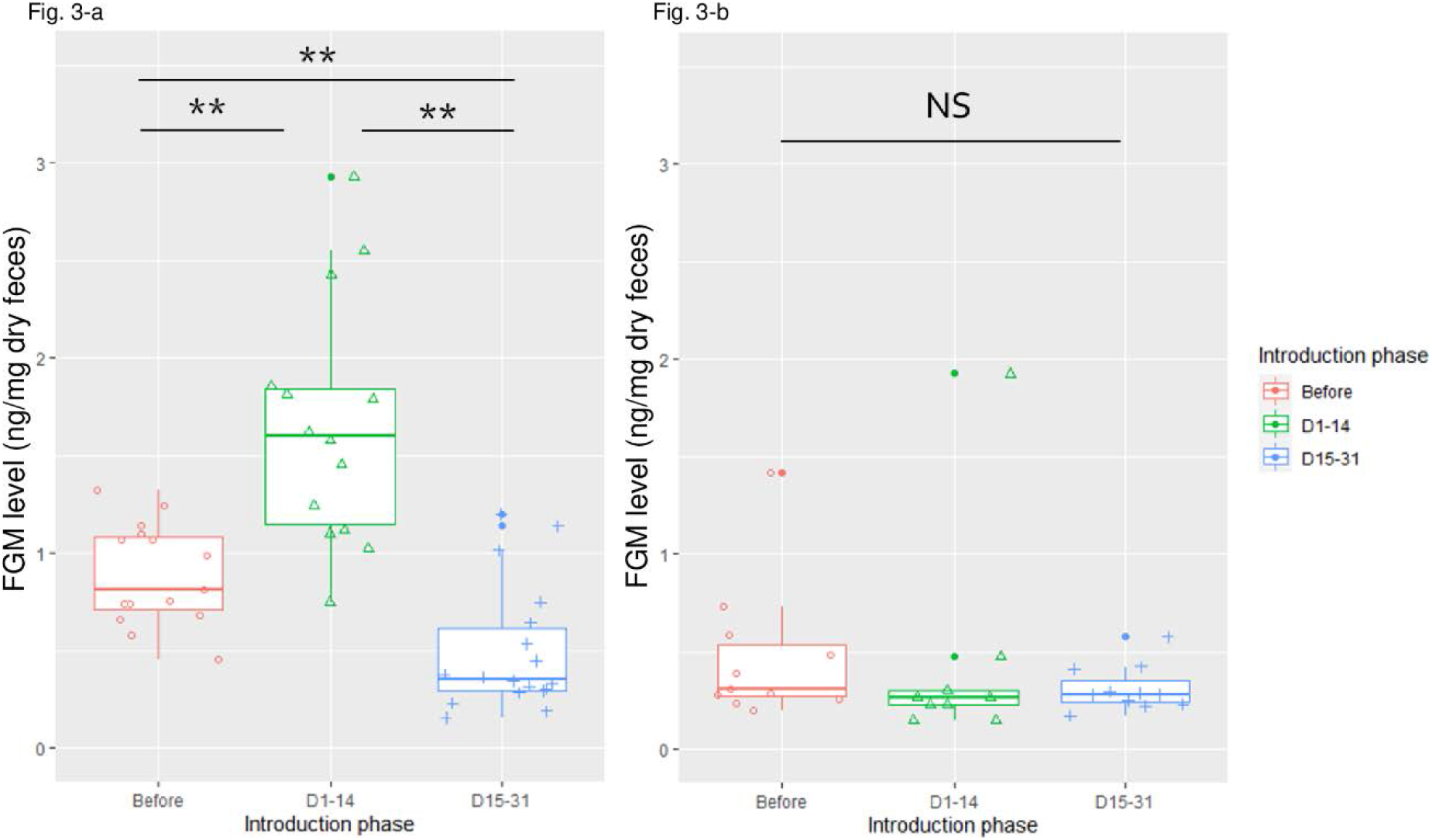
Changes in FGM levels before and after social housing. Figure 3-a. Nagi–Poplar Figure 3-b. Olive–Kunugi

### 4. Sex differences in forming social relationships

The sex of each animal did make a difference in social behavior during the beginning phase of the social housing (Figure 4). Only male–male pairs showed aggressive behaviors, while the other types of pairs did not show any aggressive behaviors (Figure 4-a). The final model of aggressive behaviors included sex, introduction phase and the interaction of sex and introduction phase (Supplementary Tables 3 and 4). Parameter estimates of the final model indicated that aggression decreased one month after the start of social housing, compared with the beginning phase. Female–female and male–female pairs showed affiliative behaviors from the beginning, while male–male pairs showed such behaviors later (Figure 4-b). The final model of affiliative behaviors included sex, introduction phase, pair and the interaction of phase and sex (Supplementary Tables 5 and 6). All types of pairs showed similar affiliative behaviors by the end of the experiment. The final model explaining the factors associated with the rate of affiliative behaviors one month after the start of the social housings only included the pair factor, not sex (Supplementary Tables 7 and 8). Some female–female and male– female pairs showed sleep sites share from the beginning, while male–male pairs showed such behaviors later (Table 4).

**Table 4.** Sex differences in the proportion of sleep site sharing The data were obtained on the first 6 days (0 month), then again for 6 days one month after the start of the social housing (1 month). The data of final phase of male-male pairs derived from the study 1 (5 pairs). The data of female-female and male-female pairs were the average of data obtained one month after the start of the social housing (2 pairs each).

| Phase | Pair | Male-Male | Female-Female | Male-Female |
| --- | --- | --- | --- | --- |
| 0 month | Pair 1 | 0.00 | 0.17 | 0.00 |
|  | Pair 2 | 0.00 | 0.50 | 1.00 |
| 1 month | Pair 1 | 0.67 | 0.50 | 1.00 |
|  | Pair 2 | 0.00 | 0.00 | 1.00 |
| Final | Average of the pairs | 0.92 | 0.25 | 1.00 |

**Figure 4.**
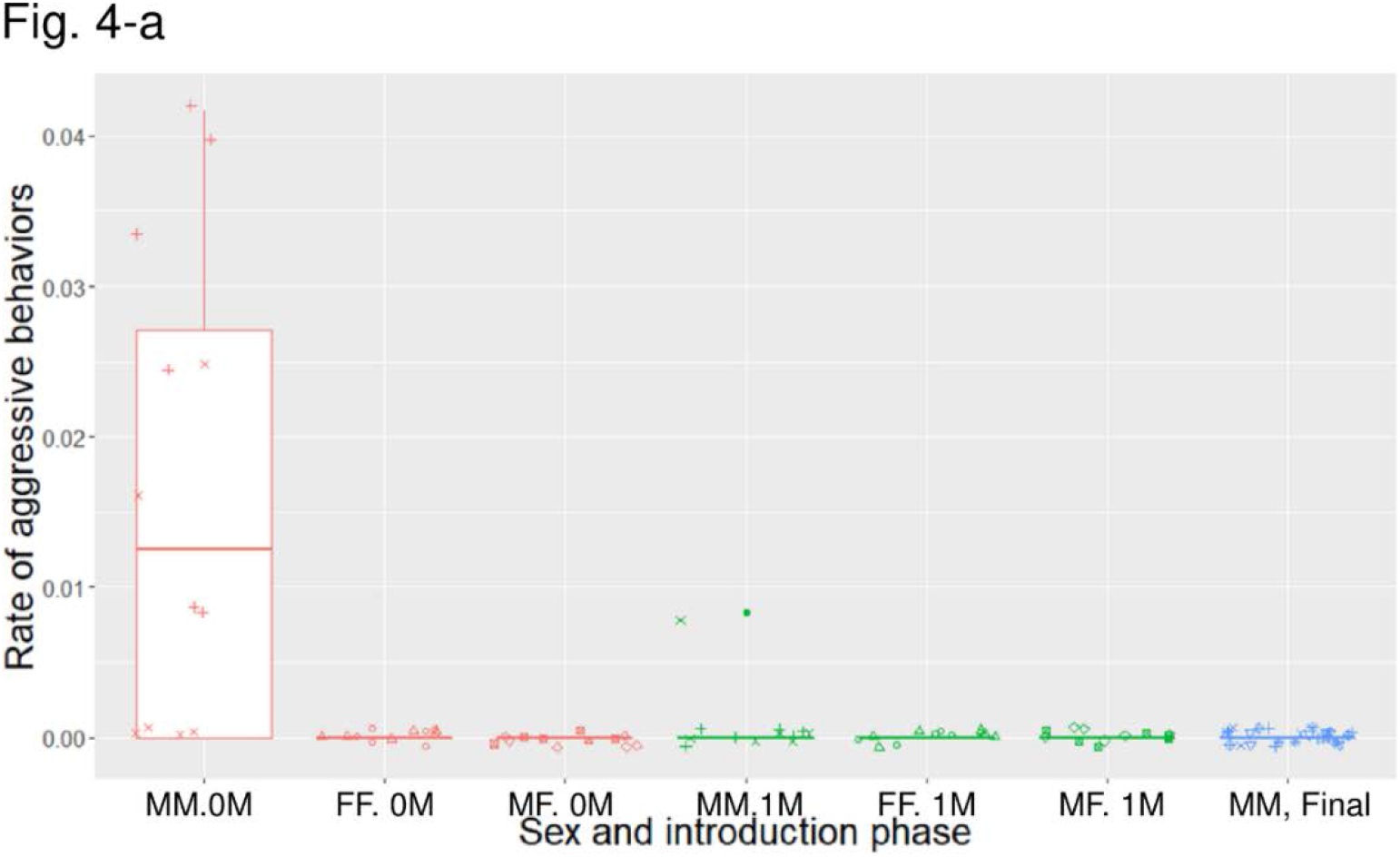
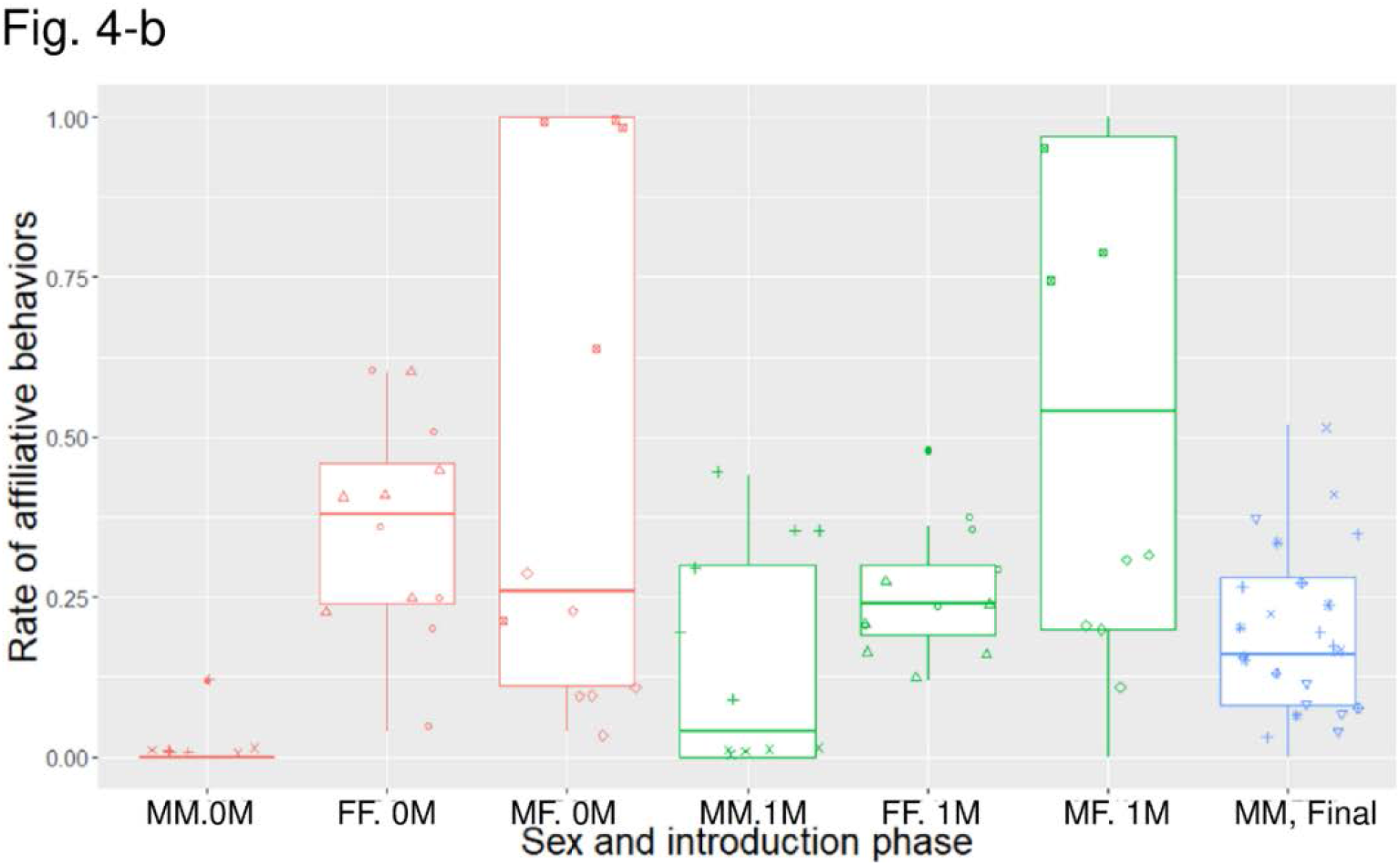
Sex differences in social behaviors during the beginning phase of social housing. FF represents data from the two female-female pairs, MF represents data from the male– female pairs, and MM represents data from the male–male pairs. The data were obtained on the first 6 days (0M), then again for 6 days one month after the start of the social housing (1M). The data of MM final derived from the study 1. Figure 4-a. Sex differences in aggressive behaviors Figure 4-b. Sex differences in affiliative behaviors

Injuries were rarely observed, with a total of six incidents requiring veterinary interventions between 2016 and 2019. Four individuals out of 16 received some form of injury (three males and one female). Most injuries were treated without separation, but the pairs in two cases were separated because the wounds did not heal well. The potential effects of social grooming on slow recovery were suspected (Supplementary table 2).

### 5. Stability of social relationship

Social relationship stability differed across the pairs. One pair showed similar levels of social behaviors between the two observation periods, another pair increased the amount of affiliative behaviors and one pair decreased the amount of affiliative behaviors and increased aggressive behaviors (Table 5). During the periods when aggressive behaviors were observed, without any affiliative behaviors, sleep sites were not shared during the light phase.

**Table 5.** Changes in social behaviors in the three male pairs between two periods

|  | Shuro – Yanagi |  |  | Nagi-Poplar |  |  | Olive-Kunugi |  |  |
| --- | --- | --- | --- | --- | --- | --- | --- | --- | --- |
| Sampling period (months) | 4 | 8 |  | 4 | 8 |  | 3 | 12 |  |
| Aggression | 0 | 1.2 ± 1.3 | W = 6, P = 0.061 | 0 | 0 | W = 18, P = 1 | 0.17 ± 0.37 | 0 | W = 21, P = 1 |
| Affiliation | 0.15 ± 0.048 | 0 | W = 36, P = 0.0022 | 0.28 ± 0.13 | 0.26 ± 0.064 | W = 19, P = 0.98 | 0 | 0.29 ± 0.19 | W = 0, P = 0.0022 |
| Sleeping site share(days) | 6 | 0 | X<sup>2</sup>=3.4, P = 0.066 | 4 | 5 | X<sup>2</sup>=0, P = 1 | 0 | 4 | X<sup>2</sup>=8.3, P < 0.001 |

## Discussion

All five pairs of male slow lorises formed affiliative relationships by the end of the first observation period. Allogrooming, social play, nest box sharing during the dark phase, and sleep site sharing during the light phase were the main behaviors observed. Some pairs ate together at the same feeding platform in close proximity at times, even if there were other places to forage. The rate of affiliative social behaviors was high in the father-son pair, but some pairs also showed comparable levels of such behaviors. One pair (Olive–Kunugi) did not show any aggressive or affiliative behaviors at times when the other pairs would have; however, even this pair showed affiliative behaviors without any aggressive behaviors by the end of the first observation period. This is surprising considering reports of the wild lorises who rarely interact with same-sex adults, but they can express social flexibility (Starr & Nekaris, 2020). In an environment without resource competition or mating pressure, males can coexist, even into adulthood. It is imperative that enclosures have adequate space and options to separate or hide from the group in captivity (Global Federation of Animal Sanctuaries, 2013), especially during the introductory phase. Additionally, it is necessary to closely monitor their relationships considering the fact that the quality of relationship of the Yanagi–Shuro pair deteriorated later.

All pairs shared sleep sites during the light phase. Sleep site sharing is often observed in various wild, nocturnal strepsirrhines (Bearder et al., 2003; Nekaris & Bearder, 2007; Rode et al., 2013), as researchers often analyze grouping tendencies by observing daytime sleep sites in the wild (Bearder et al., 2003). Although it is rare for multiple males to share sleep sites, it has been reported in the slender loris and in giant mouse lemurs (Nekaris, 2003; Rode et al., 2013). This result is not consistent with those observed in the solitary sleeping patterns of wild pygmy slow lorises (Starr & Nekaris, 2020). Given their small body size, sleeping together may have a thermoregulatory advantage at lower temperatures, but room temperature in this experiment was set within the range recommended by the husbandry manual (Fitch-Snyder et al., 2001), and there was no evidence of increased sharing at lower room temperatures. On the contrary, one pair (Tsuge–Poplar) tended to increase their sharing rate when the temperature increased, which was not expected. Consideration has to be made for the fact that they rarely slept alone, and for the temperature range, which leads us to conclude that the trend was not strong. Furthermore, all the pairs used multiple sleep sites together, and the rate of sleep site sharing was above the chance level for all pairs. These results suggest that the pygmy slow lorises in this study preferred to stay together with other individuals during the light phase. In addition, relationships with both aggressive and affiliative behaviors during the dark phase suggest that sleep site sharing could only be observed during periods when their relationship was more affiliative. Therefore, sleep site sharing is a simple and good indicator of their relationship status.

The two pairs of males that showed aggression at the beginning of the experiment, had stopped displaying aggressive behaviors around day 10. Most aggressive behaviors did not appear to be serious, and veterinary interventions were rare. We did not record any deaths that were suspected to be related to the wounds incurred through aggressions, but sometimes, recovery from injuries took time. There are previous studies that have reported a few cases of death that may have been related to trauma or infection secondary to bite wounds (Fuller, Lukas, Kuhar, & Dennis, 2014). In particular, slow lorises are venomous primates (Nekaris et al., 2013), which results in a need for close monitoring whenever adult pairs are formed to avoid serious injury or death.

Results of the changes in FGM levels for Nagi–Poplar indicates that their FGM levels increased during their phase of heightened aggression, and decreased in the later phase. The final FGM levels were even lower than those in the period when they were housed alone in the small cage or an enclosure. This result suggests that forming pairs in a more enriched enclosure is beneficial to reducing the level of physiological stress. This interpretation of these benefits was supported by the fact that the FGM levels of Nagi before social housing were higher than those of Nagi collected in the same period. However, the results from Olive–Kunugi were not consistent with that of Nagi–Poplar. There was no significant change in the level of FGM for Olive–Kunugi, which is counter intuitive. This might be related to the fact that Olive and Kunugi already lived in an enriched environment prior to the start of this study. Therefore, in addition to the heightened aggression, changes in the physical environment may have affected the increase in FGM levels of the Nagi-Poplar pair. Nevertheless, combined with the behavioral results and FGM levels, there was no evidence of increased stress over a long period of social housing. Instead, forming same-sex pairs increased the opportunity for affiliative social behaviors.

There were sex-dependent differences in the formation process of social groups. Aggressive behaviors were observed in the male–male pairs at the start of the experiment. The male–female and female–female pairs showed affiliative behaviors from the beginning. This low level of aggressive behaviors between females is interesting, considering the fact that both adult males and females in the wild do not overlap their home range with same-sex individuals (Starr & Nekaris, 2020). The details in the behaviors seen in female–female pairings will be reported in another paper (Alejandro et al. in prep). Injury risk was relatively high in the male–male pairs, but an injury was observed in a female-female pair once. Nevertheless, all types of pairs showed affiliative behaviors by the end of the experiment, without any serious incidence of aggression.

All-male groups have been formed in several captive primate species, including gorillas (Stoinski, Kuhar, et al., 2004), chimpanzees (Morimura, Idani, & Matsuzawa, 2010; Yamanashi, Teramoto, Morimura, Nogami, & Hirata, 2018), proboscis monkeys (Sha et al., 2013) and white crowned mangabeys (Fàbregas & Guillén-Salazar, 2007). All of these species are gregarious, and some of these species also have naturally occurring all-male groups in the wild, the type of species that typically form polygynous groups [e.g. gorillas (Gatti, Levréro, Ménard, & Gautier-Hion, 2004) and proboscis monkeys (Murai, 2004)]. When mating competition is removed from the environment, as in all-male captive groups, coexistence of multiple adult males is more feasible. Fundamentally, male bonding is less easily occurred compared with females because males tend to compete for mating, which cannot be shared easily, in addition to food resources (Van Schaik, 1994). In some species, particularly male philopatric species, social bonding among males might be advantageous for mating success (Van Schaik, 1994). Pygmy slow lorises are not such species, rather they are territorial, and rarely overlap their home ranges with other same-sex adult individuals (Starr & Nekaris, 2020; Wiens & Zitzmann, 2003). Additionally, it is normal for other species to exhibit aggression, even after they establish their social relationships (Leeds, Boyer, Ross, & Lukas, 2015), which has led to the recommendation for some species to form social groups at an early age (Sha et al., 2013; Stoinski, Lukas, et al., 2004). Surprisingly, in the case of male pygmy slow lorises, aggressive behaviors were very rare once they formed affiliative relationships, even when they were introduced after sexual maturation. Familiarity could not explain these relationships, except for the father-son pair as they were housed solitary at least since 2013. We are still not sure what the proximate and ultimate mechanisms of this flexible trait are, but the results of this study indicate that this species has a high motivation to interact with conspecifics if conditions are appropriate.

We are not sure if the results of this study can be generalized to other *Nycticebus* species. Although similarity exists across different species of lorises, pygmy slow lorises are unique in terms of their body size (smallest) and high rate of twin births (Fitch-Snyder et al., 2001). It is noteworthy that the chemical compounds secreted from the brachial glands of pygmy slow lorises were less than half of those of Bengal slow lorises (Hagey, Fry, & Fitch-Snyder, 2007). Because the secretion from brachial glands is the main component of the venom, bites from pygmy slow lorises might be less venomous and thus, less risky for social housings. Nevertheless, recent findings on high sociability of other slow lorises species show the importance of social management of *Nycticebus spp*. Therefore, more studies on social behaviors of both wild and captive *Nycticebus spp*. are needed to find out better social management strategies. It might be also important to study other nocturnal strepsirhines that were also often misunderstood as solitary because of the lack of understanding of their sociality.

There were some limitations to our study pertaining to the understanding of social behaviors, and optimal social management regimes. First, we could not record vocalizations and social behaviors that did not involve direct social contact. Studies have reported that vocalizations often accompany social behaviors (Daschbach, Schein, & Haines, 1981; Zimmermann, 1985). A recent study in Javan slow loris reported that the lorises used ultrasonic vocalizations for communication (Geerah, O’Hagan, Wirdateti, & Nekaris, 2019), and reported that lorises emitted vocalizations in the context of following or leading each other, and entering a sleep site. In this study, we sometimes observed that the pygmy slow lorises looked into a nest box where another individual was already inside, or kept in social proximity for a while in front of the nest box before entering the box together. It would be interesting to explore the details of pygmy slow loris communication prior to social behavior initiation in future studies. A second limitation is the fact that the judgement criteria used to test pair compatibility during the first introduction trial was obscure. We introduced two unfamiliar individuals based on published suggestions to minimize aggression during the introduction of a male and female pair (Fitch-Snyder et al., 2001). We judged their compatibility based on behaviors displayed during their first encounters. One male always showed aggressive tendencies to all individuals introduced, thus, we did not form any long-lasting pair with him. We are not sure whether his aggression was a personality trait or the result of some past experience. Since all other pairs managed to form affiliative relationships by the end of the study, it is possible that this individual may have successfully formed an affiliative relationship if given more time or tried with a different individual.

In conclusion, male slow lorises appeared to be motivated to interact with conspecifics. All pairs formed affiliative relationship by the end of the study, and they all preferred to sleep together during their light phases. Furthermore, there was no evidence of increasing physiological stress levels over a long period of time. These results suggest that living with conspecifics increases the behavioral options of pygmy slow lorises, and enables facilities to more efficiently use their available space and improve animal well-being. We believe that providing opportunities to interact with conspecifics is one of the most important parts of captive care for this species. Forming same-sex pairs might be a feasible strategy for their social management. Although adult male–male pairings might not be the first choice considering the risk of aggression during the introductory phase, such flexibility in social housing should be considered to reduce the number of individuals housed under suboptimal conditions in sanctuaries, zoos and rehabilitation centers.

## Acknowledgement

This study was financially supported by captive care grant of the International Primatological Society, Ishizue of Kyoto University, and Japan Society for Promotion of Science (17K17828, JSPS-LGP-U04, JSPS core-to-core CCSN). It is also supported by the Collaborative Research Program of Wildlife Research Center, Kyoto University. We thank Tetsuro Matsuzawa, Genichi Idani, Naoto Kimura, Koshiro Watanuki, Mari Hirosawa, Rui Hirokawa, Makiko Uchikoshi, Yuko Tawa, Yuta Shintaku and staff of the Japan Monkey Center for their support for this project. We also thank Kei Matsushima, Shiro Koshima, Kodzue Kinoshita, Satoshi Hirata, Takashi Hayakawa, Shinichi Kioka for their valuable advice and kind cooperation for this study. Thanks are also due to Anna Nekaris for her valuable advice for this project. We thank the editor and two anonymous reviewers for their constructive comments. We have no conflict of interest to declare.

---

**Supplementary Table 1.** Subject information

| Name | Sex | Date of arrival | Origin | Pairing |
| --- | --- | --- | --- | --- |
| Yanagi | M | 2006/12/7 | Confiscation (Narita) | Shuro |
| Tsubaki | F | 2007/8/2 | Confiscation (Osaka) | Tsuge |
| Mizuki | F | 2007/8/2 | Confiscation (Osaka) | Nanten |
| Momiji | F | 2007/8/2 | Confiscation (Osaka) | Poplar |
| Olive | M | 2007/8/28 | Confiscation (Osaka) | Kunugi |
| Nagi | M | 2007/8/28 | Confiscation (Osaka) | Poplar |
| Kunugi | M | 2007/8/28 | Confiscation (Osaka) | Olive |
| Nanten | F | 2007/8/28 | Confiscation (Osaka) | Mizuki |
| Hiiragi | M | 2007/8/28 | Confiscation (Osaka) | Yashi |
| Aoki | M | 2007/8/28 | Confiscation (Osaka) | Unsuccessful |
| Shuro | M | 2007/8/28 | Confiscation (Osaka) | Yanagi |
| Kaede | F | 2007/8/28 | Confiscation (Osaka) | Mimoza |
| Poplar | M | 2007/8/28 | Confiscation (Osaka) | Nagi, Tsuge, Momiji |
| Tsuge | M | 2007/8/28 | Confiscation (Osaka) | Poplar, Tsubaki |
| Mimoza | F | 2007/8/28 | Confiscation (Osaka) | Kaede |
| Yashi | M | 2010.03.28 | Born at the JMC | Hiiragi |

**Supplementary Table 2.** Duration of each pair

| Pair | Type of pair | Start | End | Reason of ending the pair |
| --- | --- | --- | --- | --- |
| Nagi - Poplar | MM | 2016/8/1 | 2017/3/31 | To treat an injury of Poplar |
| Olive - Kunugi | MM | 2016/8/14 | 2019/1/21 | To form a male-female pair |
| Yashi - Hiiragi | MM | 2016/9/16 | 2018/5/30 | Yashi passed away from sickness. |
| Shuro - Yanagi | MM | 2016/11/29 | 2017/12/12 | Shuro passed away from sickness. |
| Tsuge - Poplar | MM | 2017/5/26 | 2019/10/14 | To form a male-female pair |
| Kaede - Mimoza | FF | 2017/1/7 | 2017/7/2 | To treat an injury of Mimoza |
| Nanten - Mizuki | FF | 2017/9/12 | 2018/11/5 | To form a male-female pair |
| Poplar - Momiji | MF | 2019/10/14 | Ongoing |  |
| Tsuge - Tsubaki | MF | 2019/11/27 | Ongoing |  |

**Supplementary Table 3.** AIC table for explaining the factors associated with occurrence of aggressive behaviors

| AIC | Independent variables |
| --- | --- |
| 70.0 | Sex + Phase + Sex : Phase |
| 70.6 | Sex + Phase + Pair + Sex : Phase |
| 71.4 | Sex + Phase |
| 72.0 | Phase + Pair |
| 72.0 | Sex + Phase + Pair |

**Supplementary Table 4.** Parameter estimates for the best-fit model to explain the factors associated with occurrence of aggressive behaviors

|  | Factor | estimate | SE | z | p |
| --- | --- | --- | --- | --- | --- |
|  | Intercept | 0.51 | 0.22 | 2.3 | 0.022 |
| Sex | Female- female | -21 | 4428 | -0.0050 | 1.0 |
|  | Male - female | -21 | 4428 | -0.0050 | 1.0 |
| Phase | 1 month | -3.0 | 1.0 | -2.9 | 0.0035 |
| Sex<br>Phase | Female-female : 1month | 21 | 4487 | 0.0050 | 1.0 |
|  | Male-female : 1month | 3.0 | 6345 | 0 | 1.0 |

**Supplementary Table 5.** AIC table for explaining the factors associated with the rate of affiliative behaviors

| AIC | Independent variables |
| --- | --- |
| 329 | Sex + Phase + Pair + Sex : Phase |

**Supplementary Table 6.**
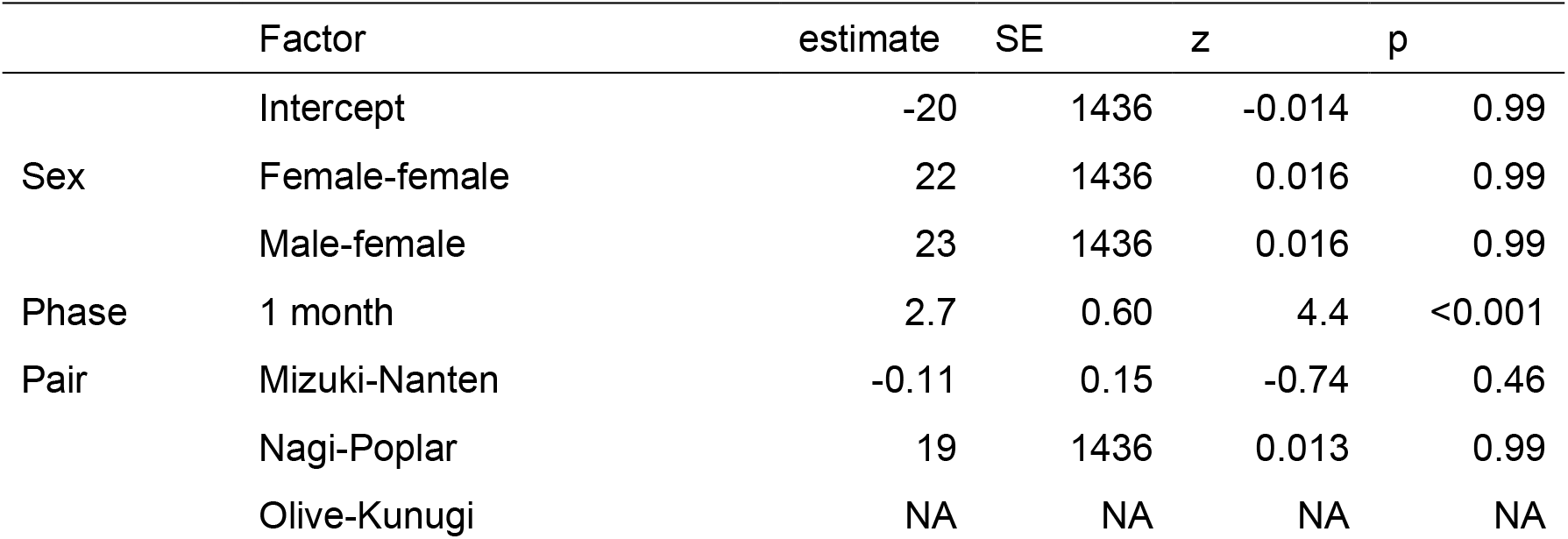
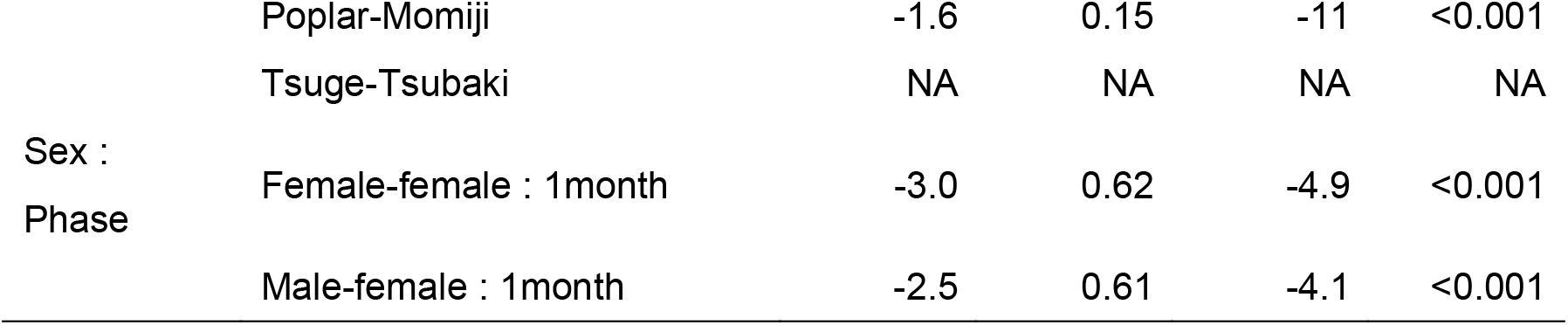
Parameter estimates for the best-fit model to explain the factors associated with the rate of affiliative behaviors

**Supplementary Table 7.**
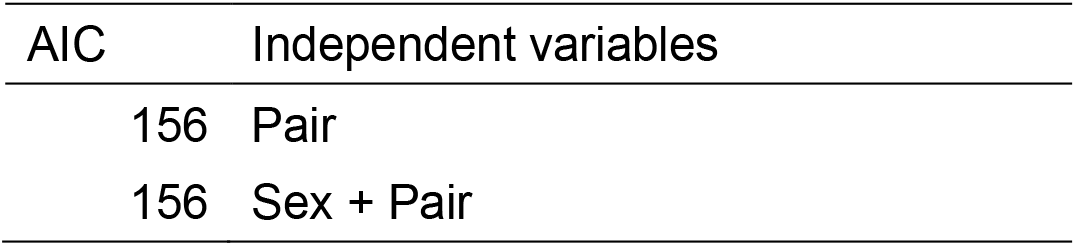
AIC table for explaining the factors associated with the rate of affiliative behaviors one month after the start of the social housings

| AIC | Independent variables |
| --- | --- |
| 156 | Pair |
| 156 | Sex + Pair |

**Supplementary Table 8.** Parameter estimates for the best-fit model to explain the factors associated with the rate of affiliative behaviors one month after the start of the social housings

|  | Factor | estimate | SE | z | p |
| --- | --- | --- | --- | --- | --- |
|  | Intercept | 2.1 | 0.14 | 14 | <0.001 |
| Pair | Mizuki-Nanten | -0.50 | 0.2 | -2.1 | 0.032 |
|  | Nagi-Poplar | -0.11 | 0.21 | -0.52 | 0.60 |
|  | Olive-Kunugi | -20 | 2334 | -0.0090 | 0.99 |
|  | Poplar-Momiji | -0.50 | 0.24 | -2.1 | 0.032 |
|  | Tsuge-Tsubaki | 1.1 | 0.17 | 6.3 | <0.001 |

**Supplementary Figure 1.**
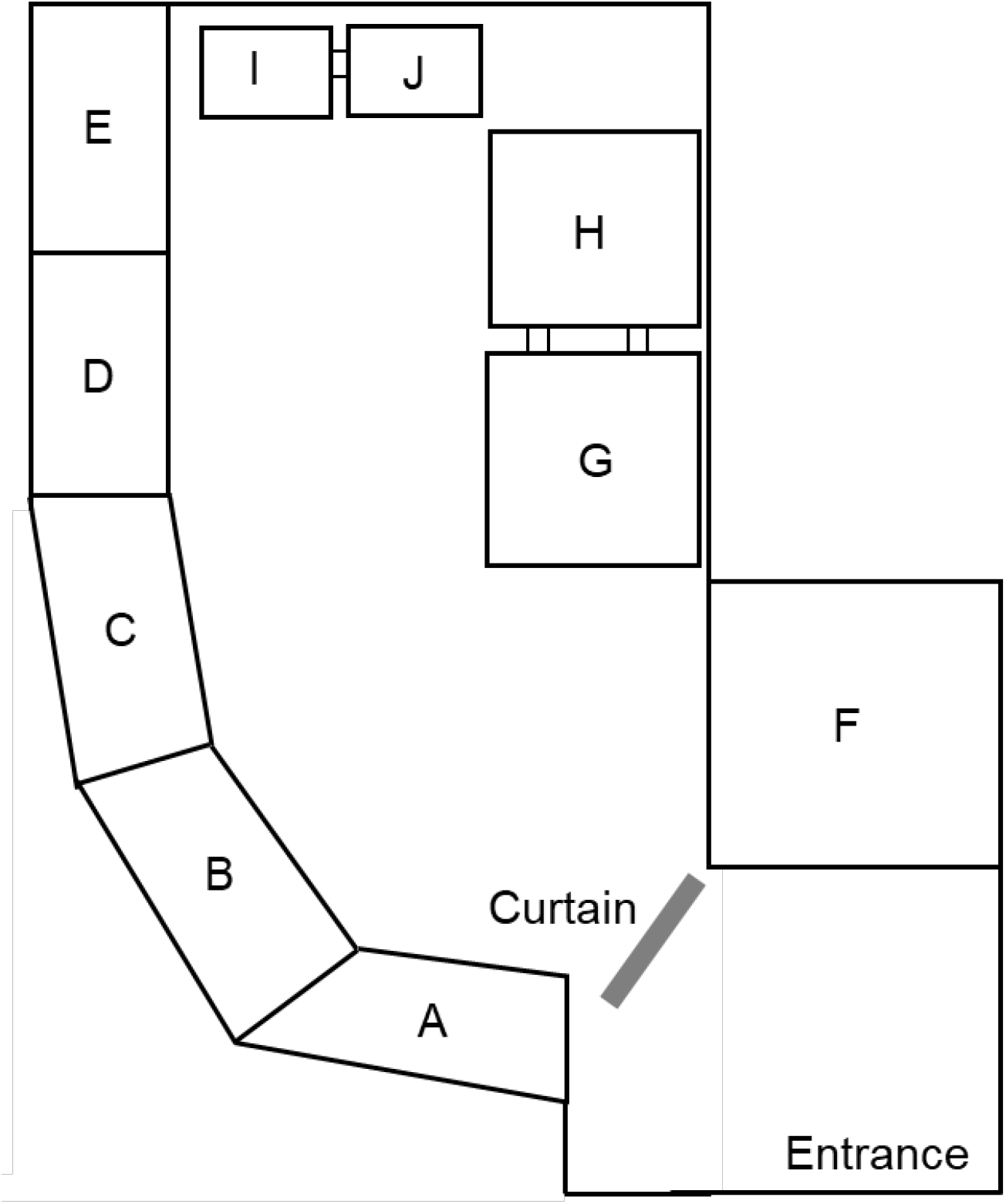
A schematic of the Slow Loris Conservation Centre (SLCC). Male-male pairings were conducted in cages A-E, while male-female and female-female pairings were conducted in all types of cages. During the study periods of male-male pairings, females lived in cages F-J. G-H and I-J can either be used as a single cage or two adjacent cages.

Supplementary video https://youtu.be/3gv1qbnppBo

